## Supplementary data for "SND3 is the membrane insertase within a fungal multipass translocon"

### Supplementary Figures and Tables

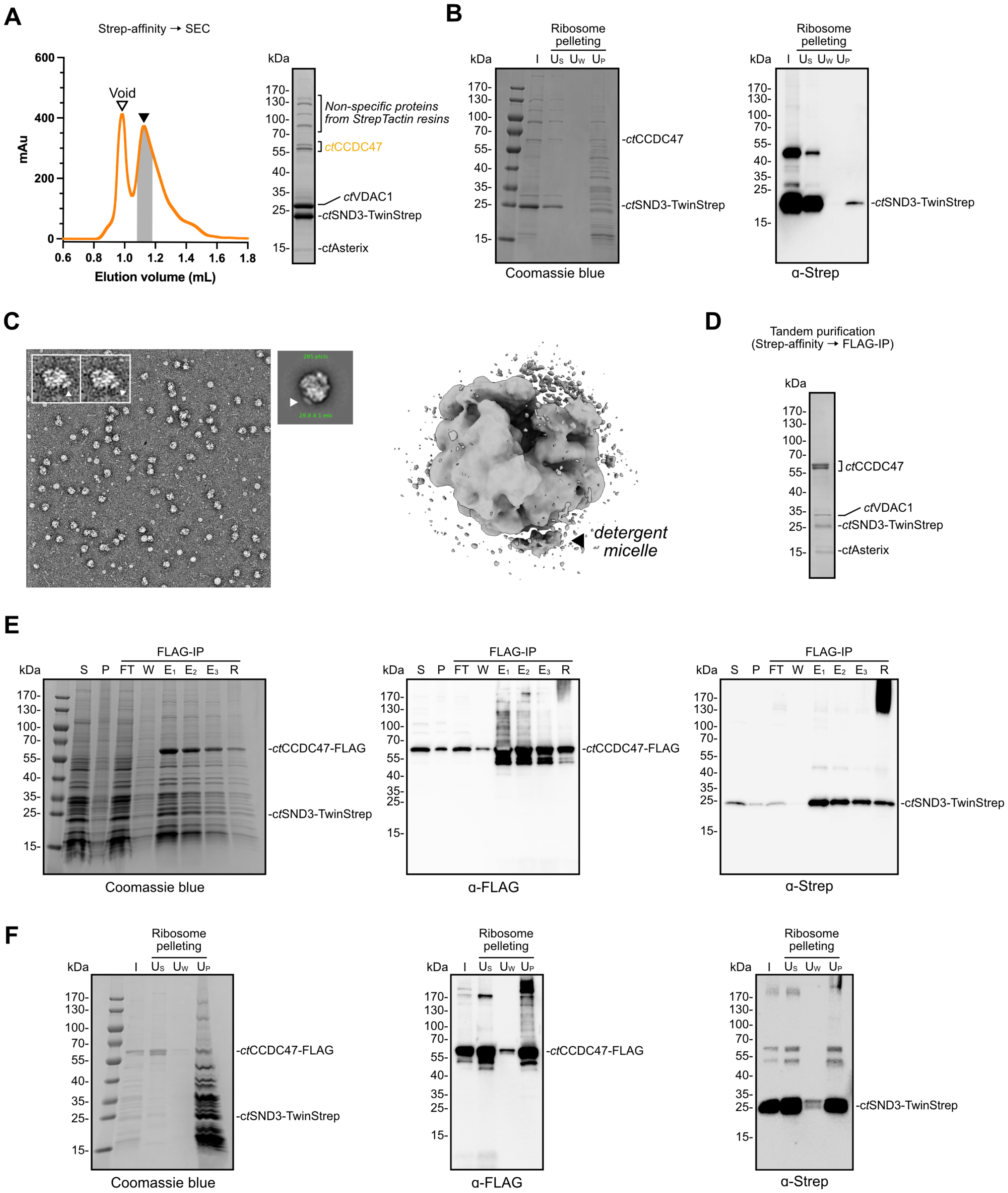


**Supplementary Fig. 1. Purification strategies for isolating ribosome-bound SND3-associated complexes.**

(**A**) Left: SEC profile of StrepTactin affinity-purified *ct*SND3-TwinStrep complexes from the *C. thermophilum* *ct*SND3-TwinStrep strain. Right: The single fraction highlighted in grey was analysed by SDS-PAGE with Coomassie blue staining. The gel bands for *ct*SND3, *ct*CCDC47 and *ct*VDAC1 were identified by MS analysis. The band for *ct*Asterix is inferred from MS analysis of the gel in **D**. (**B**) Additional ultracentrifugation after StrepTactin affinity purification of *ct*SND3-TwinStrep showed a small proportion of SND3 in the ribosome-containing pellet ($U_P$) when separated by SDS-PAGE. Samples from the affinity-purified *ct*SND3 input (I), ribosome-free supernatant ($U_S$) and pellet wash ($U_W$) are also analysed by Coomassie blue staining (left) or western blotting with monoclonal anti-Strep antibody (right). (**C**) Negative stain EM analysis of the ribosome-containing fraction in **B** showing a representative micrograph and exemplary particles (left), an example 2D class average (middle) and the resulting low-resolution map (right). Arrows in each image indicate the associated density at the ribosome tunnel exit. (**D**) SDS-PAGE analysis and Coomassie blue staining of the complex obtained after tandem StrepTactin-affinity purification and FLAG-IP from the *C. thermophilum* *ct*SND3-TwinStrep/*ct*CCDC47-FLAG strain. The labelled gel bands were identified by MS analysis. (**E-F**) Samples taken during (**E**) FLAG-IP of *ct*CCDC47-FLAG and (**F**) further ribosome pelleting were separated by SDS-PAGE and visualised by Coomassie blue staining and western blotting with anti-FLAG or anti-Strep antibodies as indicated. Aside from ribosome pelleting samples defined in **B**, samples were taken from the soluble fraction after detergent solubilisation (S), the insoluble fraction after detergent solubilisation (P), the unbound fraction from the FLAG-IP (FT), the wash fraction from the FLAG-IP (W), the three separate elution fractions from the FLAG-IP ($E_{1-3}$) and the anti-FLAG M2 affinity gel after elution (R).

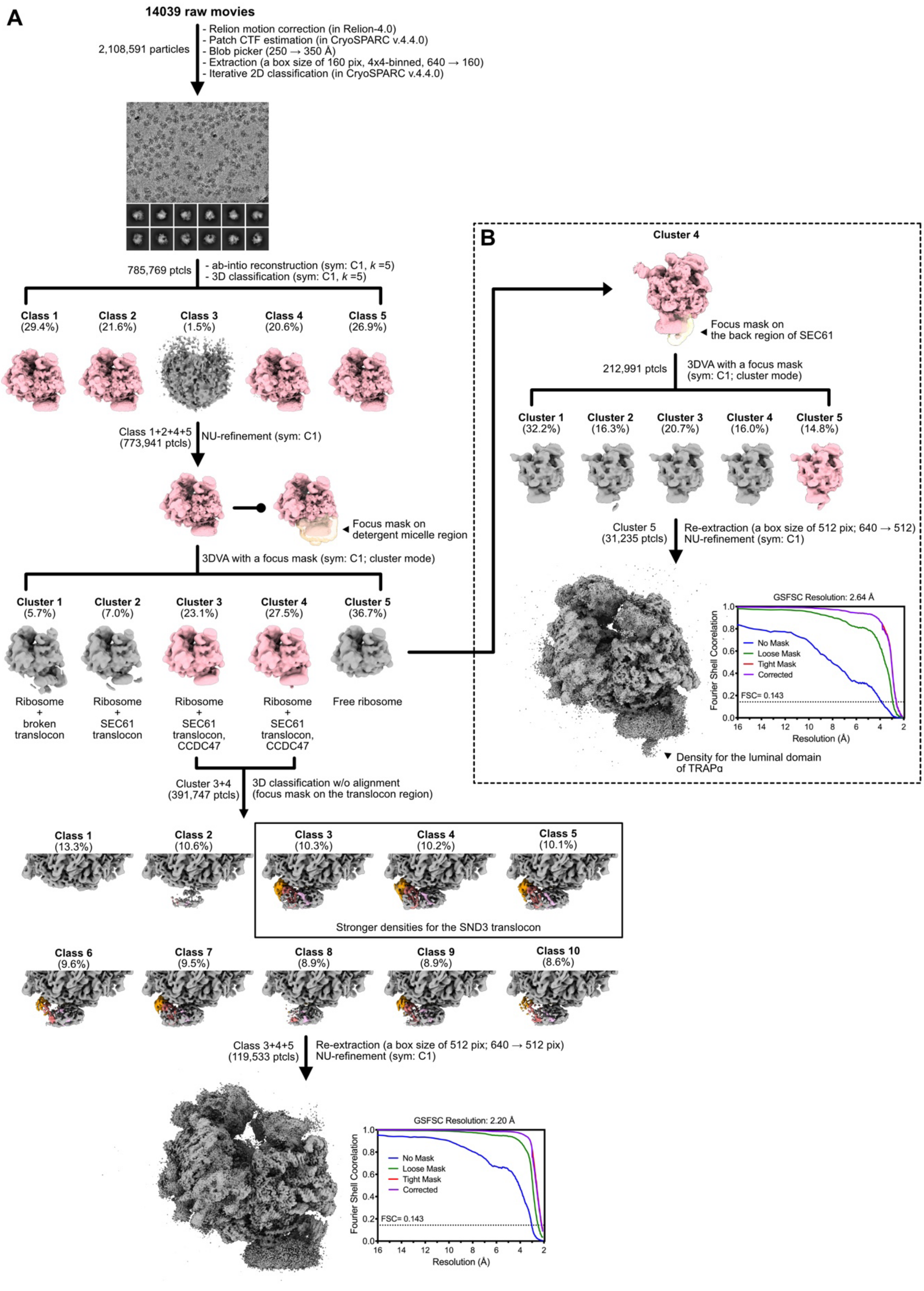


**Supplementary Fig. 2. Workflow for single-particle cryo-EM data processing of the ribosome-bound SND3 translocon.**

(**A**) Overview of the cryo-EM data processing of the ribosome-bound SND3 translocon. The micrographs that meet the selection criterion of “CTF_fit_resolution” <4 Å were used for particle selection, iterative rounds of 2D classification, 3D classification, and non-uniform (NU) refinement. The resulting cryo-EM map was used for 3D variability analysis (3DVA) with a mask focusing on the detergent micelle region. Two selected 3DVA classes were pooled and further classified by a round of focused 3D classification without alignment. The resulting three classes were selected and used to generate the final cryo-EM map of the ribosome-bound SND3 translocon at an overall resolution of 2.20 Å. (**B**) The 3DVA cluster 4 was independently selected for further processing with a focused mask on the back region of SEC61, resulting in a second cryo-EM map with improved density for the luminal domain of *ct*TRAPα. Gold standard FSC curves are shown next to the final reconstructions.

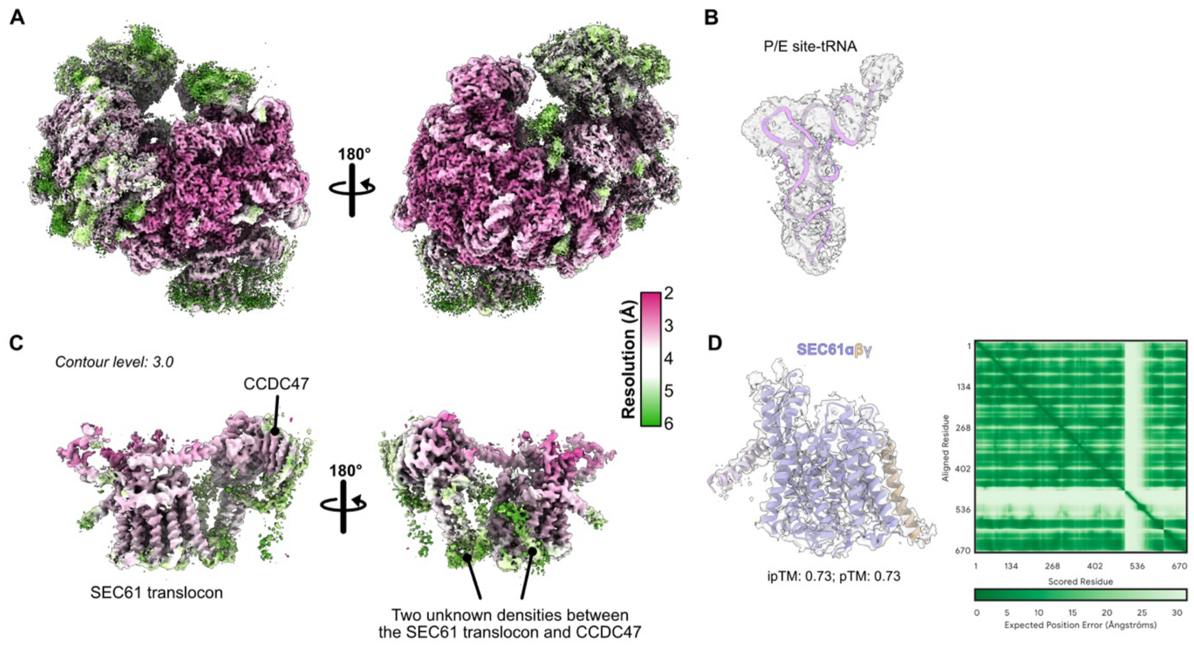


**Supplementary Fig. 3. Structural dissection and identification of the ribosome-bound SND3 translocon.**

(**A**) Local resolution estimation of the overall cryo-EM structure of the ribosome-bound SND3 translocon. (**B**) Superimposition of the final cryo-EM map with the structure of the pe/E-site tRNA from PDB 7OLD. (**C**) Zoom-in of the detergent micelle region for the views shown in A. The contour level of the cryo-EM map was set to 3.0 in UCSF-ChimeraX[56]. The schematics reveal two clear densities for the *ct*SEC61 translocon and *ct*CCDC47 and two unknown densities between the *ct*SEC61 translocon and *ct*CCDC47. (**D**) Rigid body docking of the AF3[22] model for the *ct*SEC61 translocon into the corresponding density. Confidence metrics for the AF3 model are shown, including the predicted local distance difference test (pLDDT) and predicted alignment error (PAE), as well as scores for predicted template modeling (pTM) and the interface predicted template modeling (ipTM).

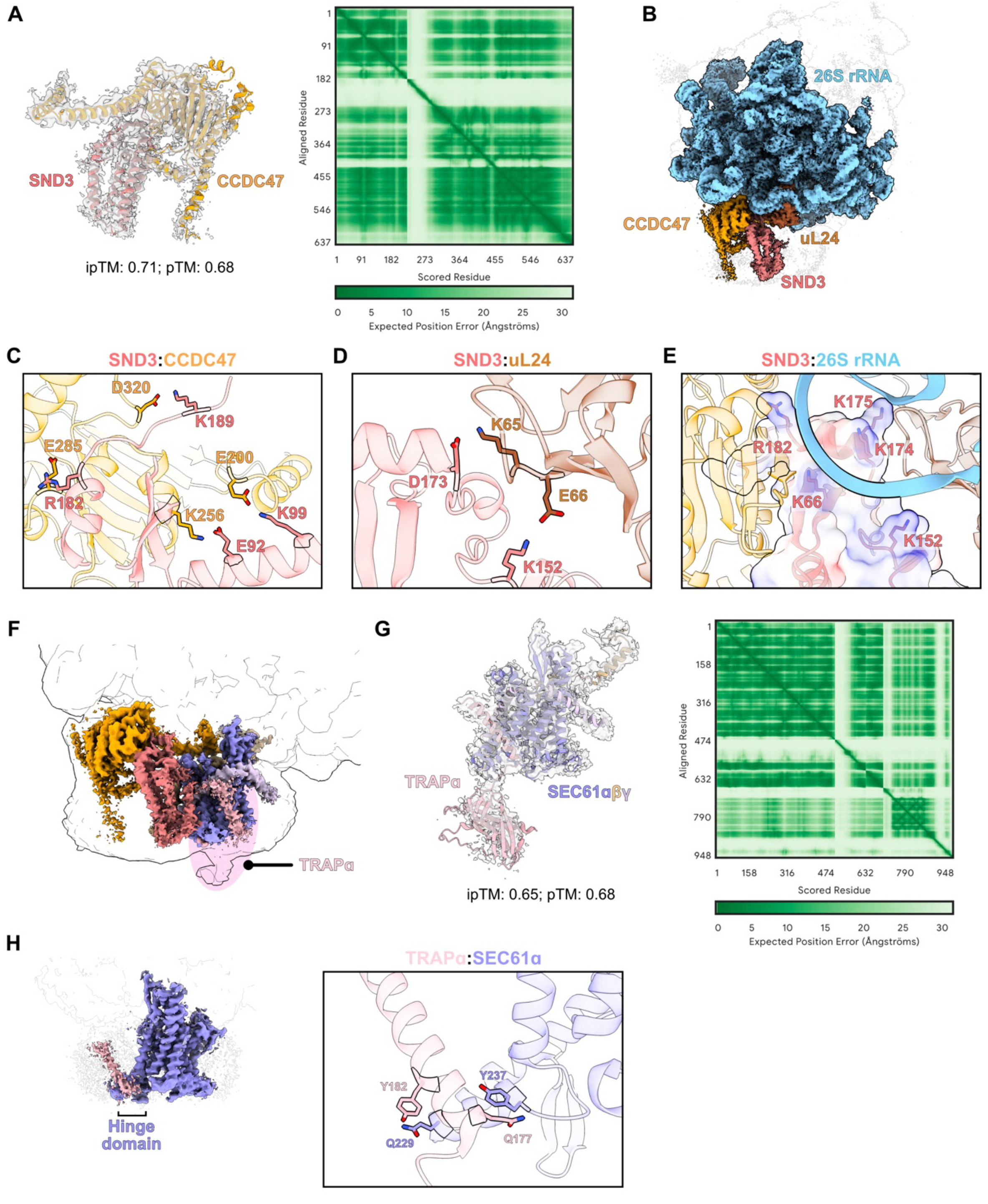


**Supplementary Fig. 4. Interactions of SND3 and TRAPα within the SND3 translocon.**

(**A**) Rigid body docking of the AF3[22] model for the *ct*CCDC47/SND3 complex into the density for *ct*CCDC47 identified the adjacent unknown density as *ct*SND3. (**B**) Overview of the interaction partners of *ct*SND3 within the SND3 translocon complex. Detailed views of the interactions of *ct*SND3 with (**C**) *ct*CCDC47, (**D**) uL24, and (**E**) 26S rRNA are shown. (**F**)

Superimposition of the final cryo-EM map and the corresponding binned map (4x binning) at low contour levels. This schematic reveals the second unknown density comprises a TMD and an ER-luminal domain, which resembles the structure of TRAPα. (**G**) Structural identification of *ct*TRAPα in the SND3 translocon by rigid body docking of an AF3 model of the *ct*SEC61 translocon/TRAPα complex into the improved cryo-EM map shown in **Supplementary Fig. 2B**. (**H**) Views of the interaction between the TMD of *ct*TRAPα and the hinge domain of *ct*SEC61α in the cryo-EM reconstruction (left) and model (right). For all AF3 models, the predicted local distance difference test (pLDDT) and predicted alignment error (PAE) are shown, as well as scores for predicted template modeling (pTM) and the interface predicted template modeling (ipTM).

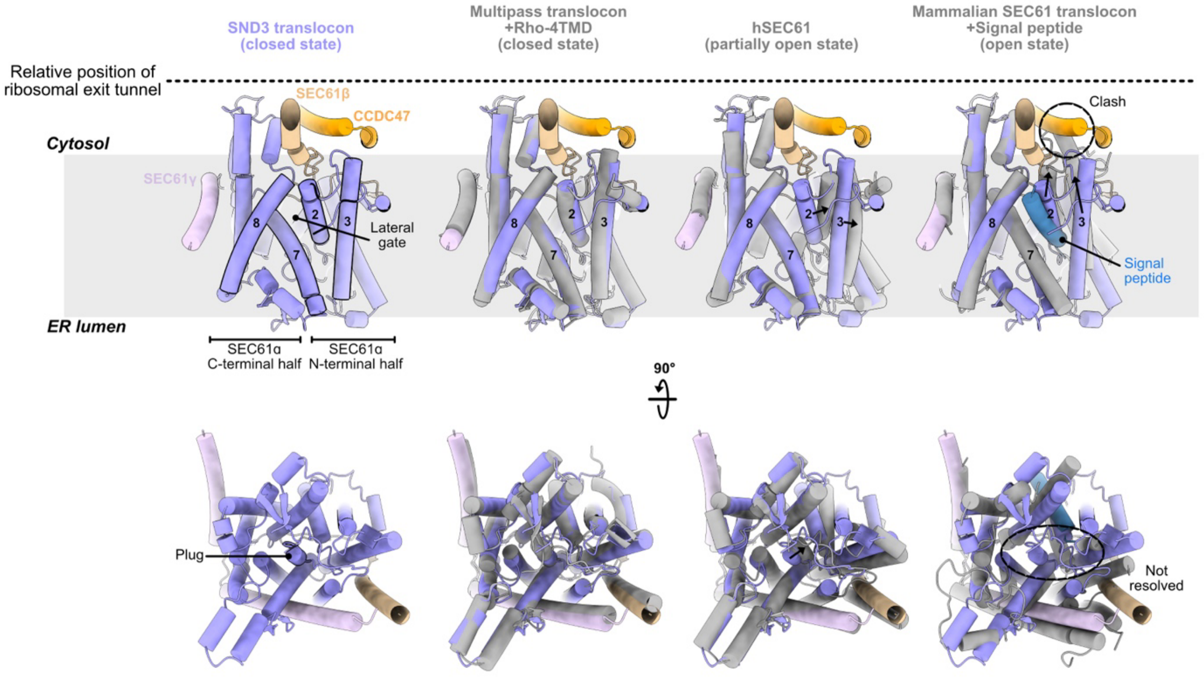


**Supplementary Fig. 5. The SEC61α channel adopts a closed conformation in the SND3 translocon.**

Side-by-side comparison between *ct*SEC61α in the SND3 translocon (purple) and representative structures of the channel in different states (grey). The C-terminal half of *ct*SEC61α is superimposed with structures for the closed (PDB 7TUT; RMSD 1.06 Å over 241 atoms), partially open (PDB 8DNV; RMSD 1.13 Å over 210 atoms) and open (PDB 3JC2; RMSD 1.01 Å over 132 atoms) SEC61α channel structures. (Top) View from the membrane plane. Arrows show the different position for TMD2 and TMD3 in the partially open and open channels, and the circle highlights a clash between *ct*CCDC47 (orange) and the cytosolic end of TMD3 in the open channel. (Bottom) View from the ER lumen. The arrow shows the movement of the plug helix in the partially open channel and the circle indicates that the plug helix is not resolved in the open channel.

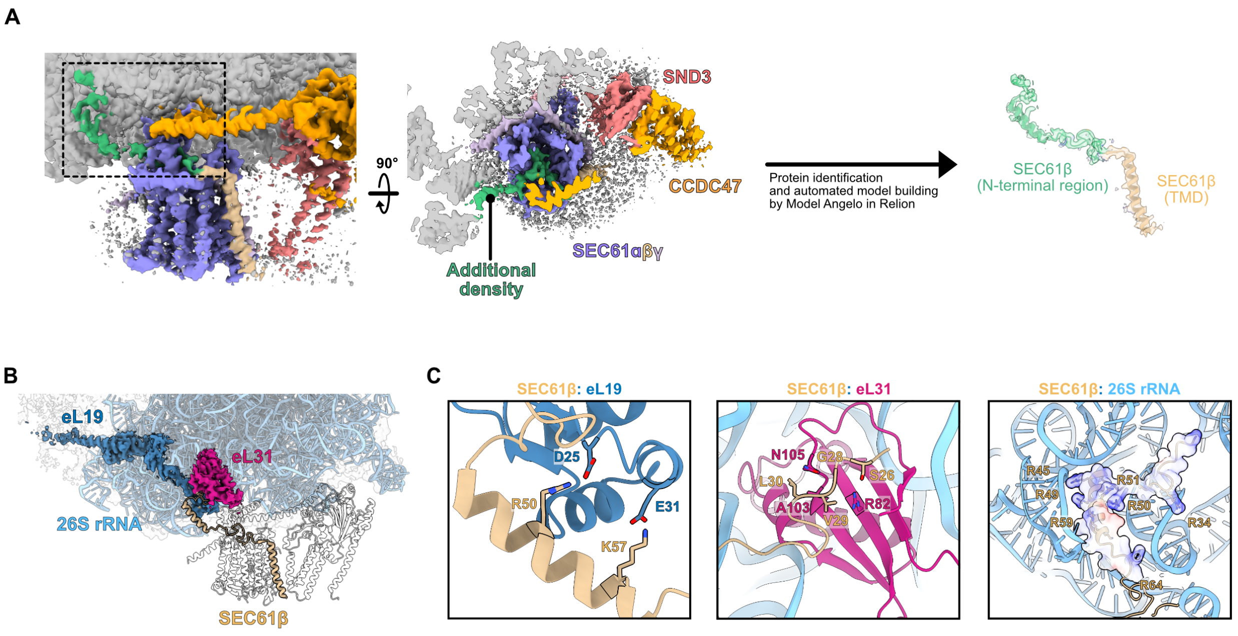


**Supplementary Fig. 6. Structural characterisation of the N-terminal region of SEC61β.**

(**A**) Additional density (green) is found attached to the ribosome and extends to the entry of the *ct*SEC61 translocon. Protein identification as the N-terminal region of *ct*SEC61β and automatic model building in the density were performed using Model Angelo[27]. (**B**) Overview of *ct*SEC61β interactions with the ribosome. (**C**) Detailed views of the interfaces between the *ct*SEC61β ribosome binding domain and eL19, eL31 and 26S rRNA.

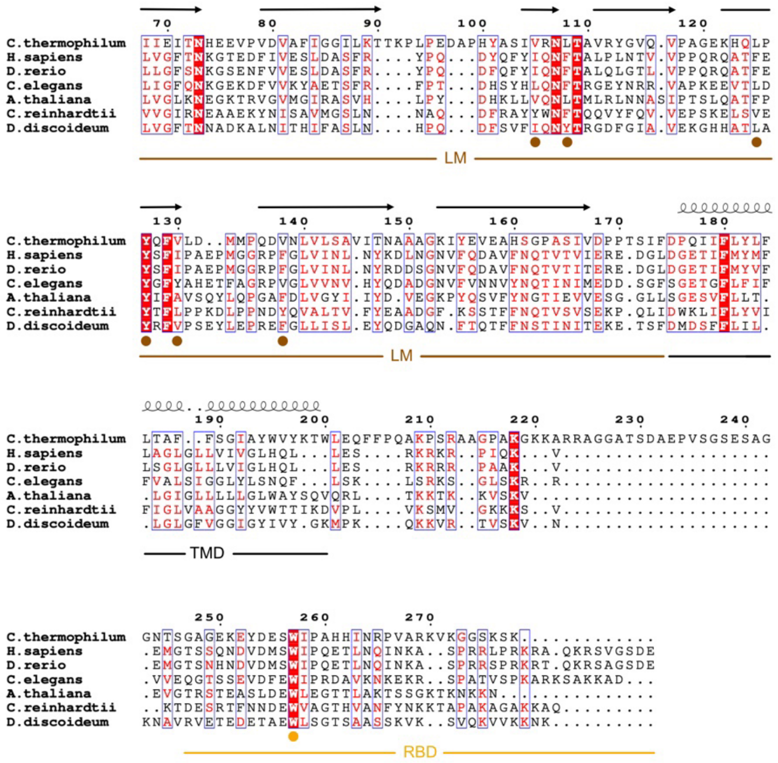

**Supplementary Fig. 7. TRAPα sequence conservation in eukaryotes.**

Sequence alignment of TRAPα homologues generated using Clustal Omega[87] and visualised using Espript 3.0[88]. Secondary structure elements from the AF3 model of *ct*TRAPα are shown above the sequence for the luminal domain (LM). Circles below the alignment highlight conserved residues shown to be functionally important in the LM[24] (brown) and for the interaction between the TRAPα ribosome binding domain (RBD) and the ribosome (orange).

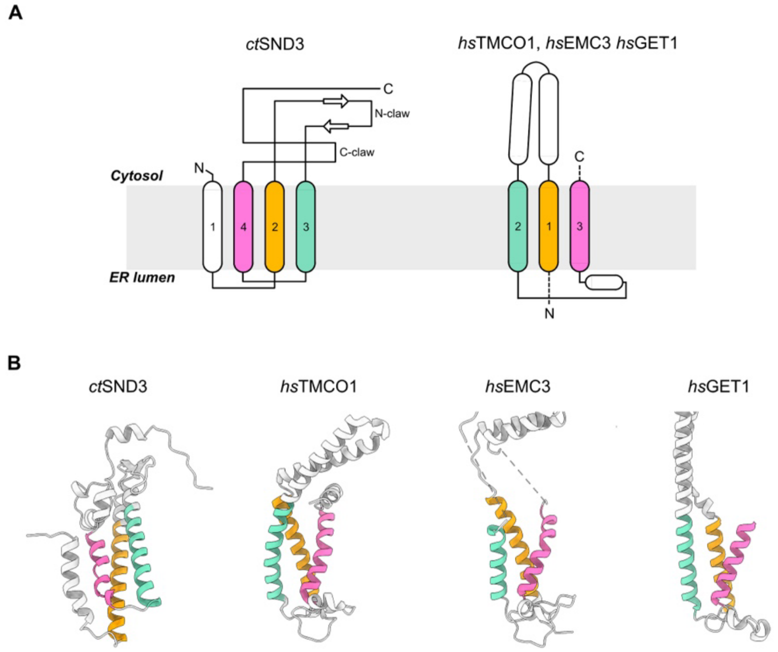


**Supplementary Fig. 8. *ct*SND3 does not have the fold of an Oxa1 superfamily insertase.**

(**A**) Schematic representation of the topology of *ct*SND3 and human (*hs*) Oxa1 superfamily membrane insertases showing a swapped TMD organization and different secondary structure of the extramembranous loops relative to *ct*SND3. The three TMDs contributing to the membrane-embedded hydrophilic groove are coloured in sequence order. (**B**) Side-by-side comparison of the structures of *ct*SND3 and human Oxa1 superfamily membrane insertases coloured as in **A**. *hs*EMC3 (PDB 6WW7) was superimposed with *hs*TMCO1 (PDB 7TUT; RMSD 1.40 Å over 30 atoms) and *hs*GET1 (PDB 8CR1; RMSD 1.17 Å over 31 atoms).

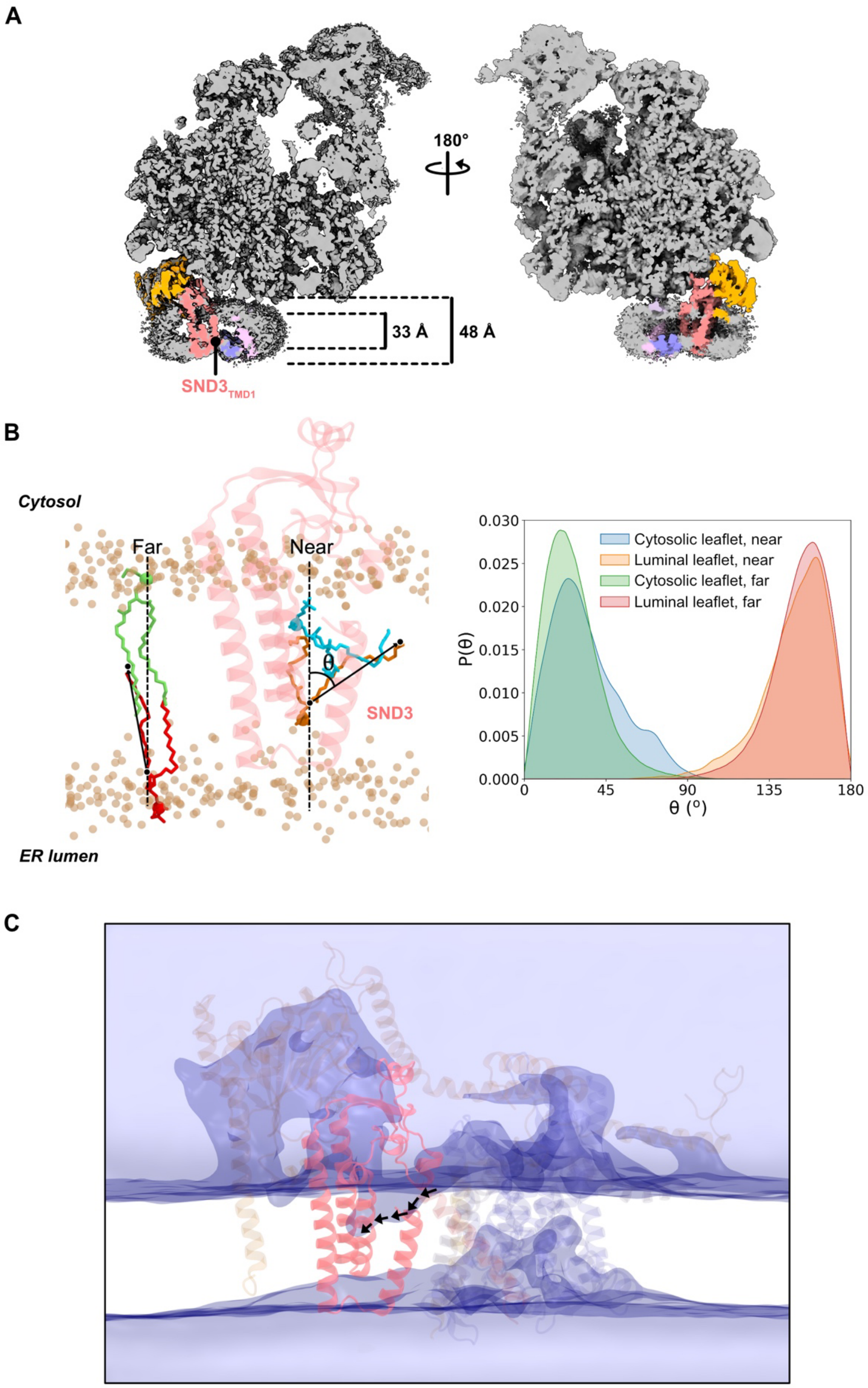


**Supplementary Fig. 9.** ***ct*****SND3 causes local membrane thinning.**

(**A**) Slices of the cryo-EM map of the SND3 translocon through the membrane normal demonstrate local detergent micelle thinning occurring in the vicinity of *ct*SND3 TMD1. The density corresponding to each SND3 translocon component is coloured as in **Fig. 1B**. (**B**) (Left) a simulation snapshot showing the membrane, where phospholipids are categorised as 'near' (within 3 Å of *ct*SND3) or 'far' (more than 10 Å away). (Right) distribution of the angles between phospholipid tails and the membrane normal. The angle (θ) is defined between the z-axis (membrane normal) and the vector connecting the first and last atoms of each hydrophobic lipid chain. These distributions reveal distortions in lipid orientation near *ct*SND3 in both the cytosolic and luminal membrane leaflets. (**C**) Snapshot of an atomistic MD simulation showing the density of water above and below the membrane as a transparent purple isosurface. The structure of the SND3 translocon is shown in cartoon representation, in which *ct*SND3 is opaque and other components are transparent. The arrows indicate a pathway for water molecules from the cytosol to the hydrophilic groove of *ct*SND3.

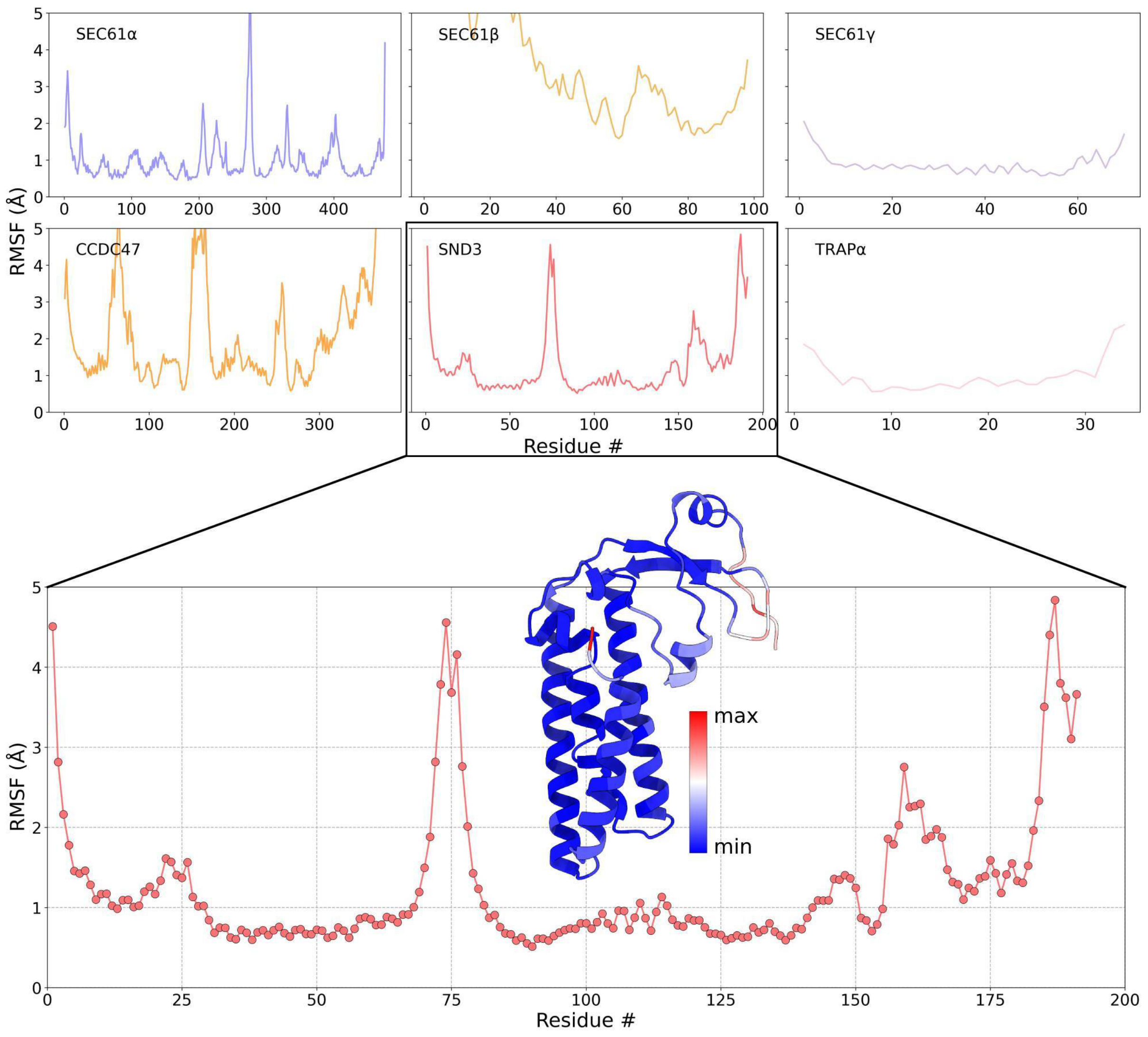


**Supplementary Fig. 10. Structural rigidity of the SND3 translocon in atomistic MD simulations.**

Rigidity of protein chains quantified by root mean square fluctuations (RMSF) of backbone Cα atoms. The top panel shows the RMSFs for all the protein chains present in the SND3 translocon complex. The RMSF of SND3 is shown separately in the bottom panel. The cartoon representation of SND3 is colored according to B-factors (directly proportional to RMSF) of the atoms. The color gradient from blue to red denotes increasing flexibility as shown in the color bar.

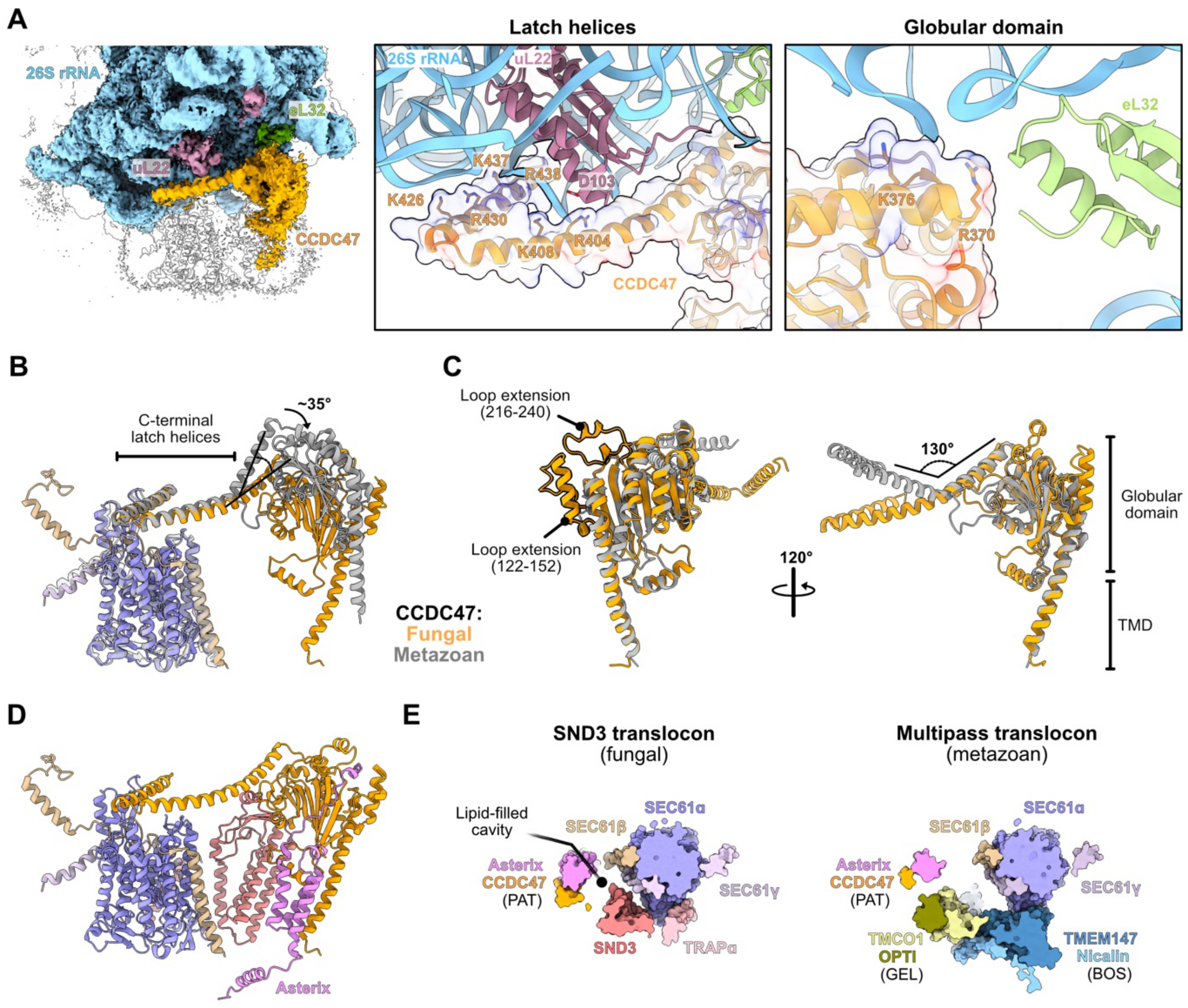


**Supplementary Fig. 11. Comparison of the PAT complex between the fungal and metazoan MPT.**

(**A**) Overview of *ct*CCDC47 interactions with the ribosome (left), showing detailed views of the interfaces with uL22, eL32 and 26S rRNA (right). The interface with the *ct*CCDC47 latch helices is conserved with the metazoan MPT, whilst the interface with the globular domain is not. (**B**) Structural comparison of the SND3 translocon (colour) and metazoan MPT (grey; PDB 7TUT), superimposed as in **Fig. 4**. The latch helices of CCDC47 are positioned identically relative to the SEC61 translocon, but the globular domains are titled by ~35º with respect to each other. (**C**) Superimposition of the TMD and globular domain of fungal (orange) and metazoan CCDC47 (grey; RMSD 1.30 Å over 170 atoms) indicates a largely similar fold aside from the two indicated helical loop extensions *ct*CCDC47. The connection to the latch helices is kinked by 130º in metazoan CCDC47 resulting in the repositioning shown in B. (**D**) Model for the interaction of *ct*Asterix with the SND3 translocon shown in the same view as **Fig. 1C**.

The position of *ct*Asterix was derived from an AF3[22] model for the *ct*Asterix/CCDC47/SND3 ternary complex that had been superimposed with *ct*CCDC47 in our structure (RMSD 0.80 Å over 294 atoms). (**E**) Comparison of the model in D with the metazoan MPT, superimposed and represented as in **Fig. 4A**.

| Protein | Complex | Fungi | Metazoa | Viridiplantae | Alveolata | Stramenopiles | Euglenozoa | Amoebozoa | |
|---|---|---|---|---|---|---|---|---|---|
| SND3 | | 2107 | 3 | 14 | 16 | 137 | 29 | 9 | fungal MPT |
| TMCO1 | GEL | 79 | 1521 | 425 | 70 | 6 | 0 | 15 | |
| OPTI | GEL | 75 | 1426 | 479 | 46 | 6 | 0 | 9 | |
| TMEM147 | BOS | 0 | 1423 | 411 | 47 | 2 | 0 | 8 | metazoan MPT |
| Nicalin | BOS | 0 | 1830 | 444 | 45 | 8 | 27 | 0 | |
| NOMO | BOS | 0 | 1690 | 423 | 5 | 5 | 0 | 11 | |
| CCDC47 | PAT | 1890 | 1655 | 525 | 68 | 33 | 1 | 15 | |
| Asterix | PAT | 1435 | 1378 | 377 | 9 | 1 | 1 | 8 | |
| SEC61α | | 2393 | 2269 | 567 | 116 | 97 | 36 | 18 | conserved components |
| SEC61β | | 2334 | 1097 | 1003 | 87 | 58 | 4 | 5 | |
| SEC61γ | | 1781 | 2864 | 27 | 70 | 73 | 24 | 15 | |
| TRAPα | | 1806 | 1783 | 616 | 1 | 9 | 29 | 10 | |

**Supplementary Fig. 12. Mutually exclusive conservation of the fungal and metazoan MPT.**

Homologues for the fungal-specific, metazoan-specific and conserved MPT components were identified in the given eukaryotic taxa using OrthoDB v12.0[86]. Each column is coloured on an independent relative scale of lowest (white) to highest (dark grey) number of homologues.

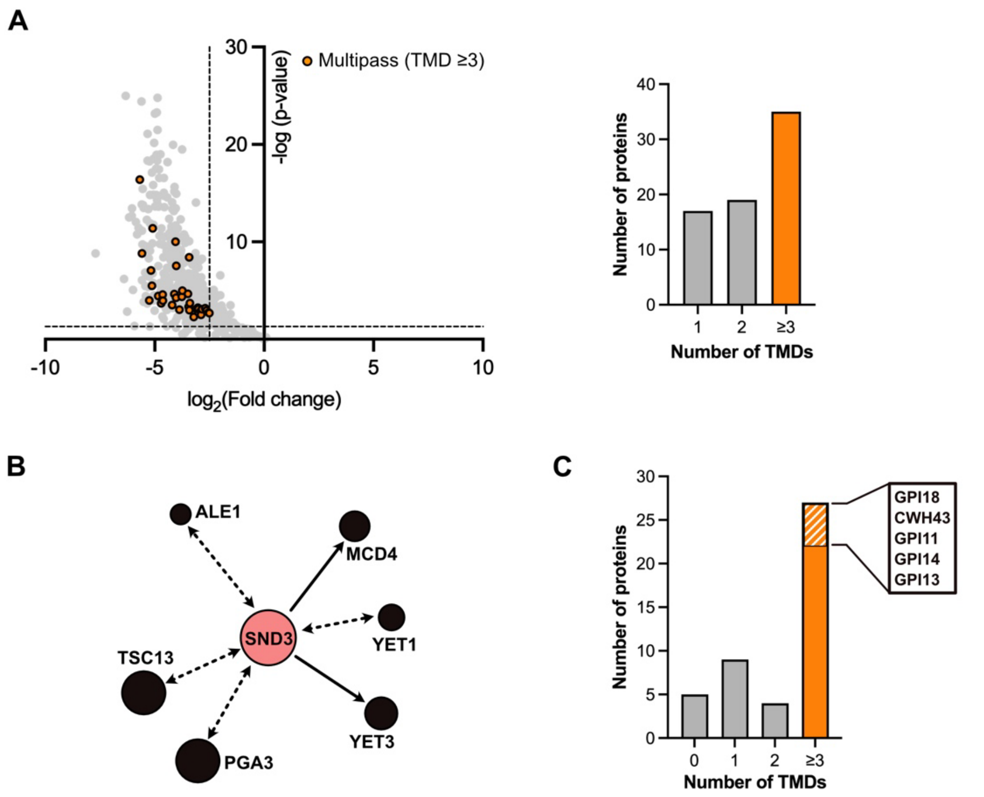


**Supplementary Fig. 13. Evidence that the SND3 translocon functions in multipass IMP biogenesis.**

(**A**) MS differential abundance analysis showing multipass IMPs are enriched after FLAG-IP from the *C. thermophilum ct*CCDC47-FLAG/*ct*SND3-TwinStrep strain. Left: Volcano plot showing proteins enriched in samples taken after FLAG-IP compared to equivalent control samples purified from wild type *C. thermophilum*. Differential abundance analysis was conducted on four independent biological purifications. Multipass IMPs with ≥3 TMDs predicted by DeepTMHMM 1.0[89] were identified and those which are significantly enriched ($log_2$(fold change) <-2.5, p-value <0.05) in our sample are highlighted in orange, excluding components of the SND3 translocon. Right: The absolute number of multipass IMPs (orange) is enriched relative to single or double-spanning IMPs. (**B**) The SND3/PHO88 cluster extracted from the yeast-interactome web application (http://www.yeast-interactome.org/)[35] comprises entirely multipass IMPs. Arrows show interactors of SND3, with dashed arrows indicating interactions only supported by profile correlation. (**C**) Proteins whose biogenesis was affected in a *S. cerevisiae Δsbh1/Δsbh2* strain[36] (SBH1 and SBH2 are the yeast homologues of SEC61β) were classified according to the number of TMDs, as predicted by DeepTMHMM 1.0.

Multipass IMPs (orange) are predominantly affected and the subset involved in GPI anchor biosynthesis are highlighted (orange stripes).

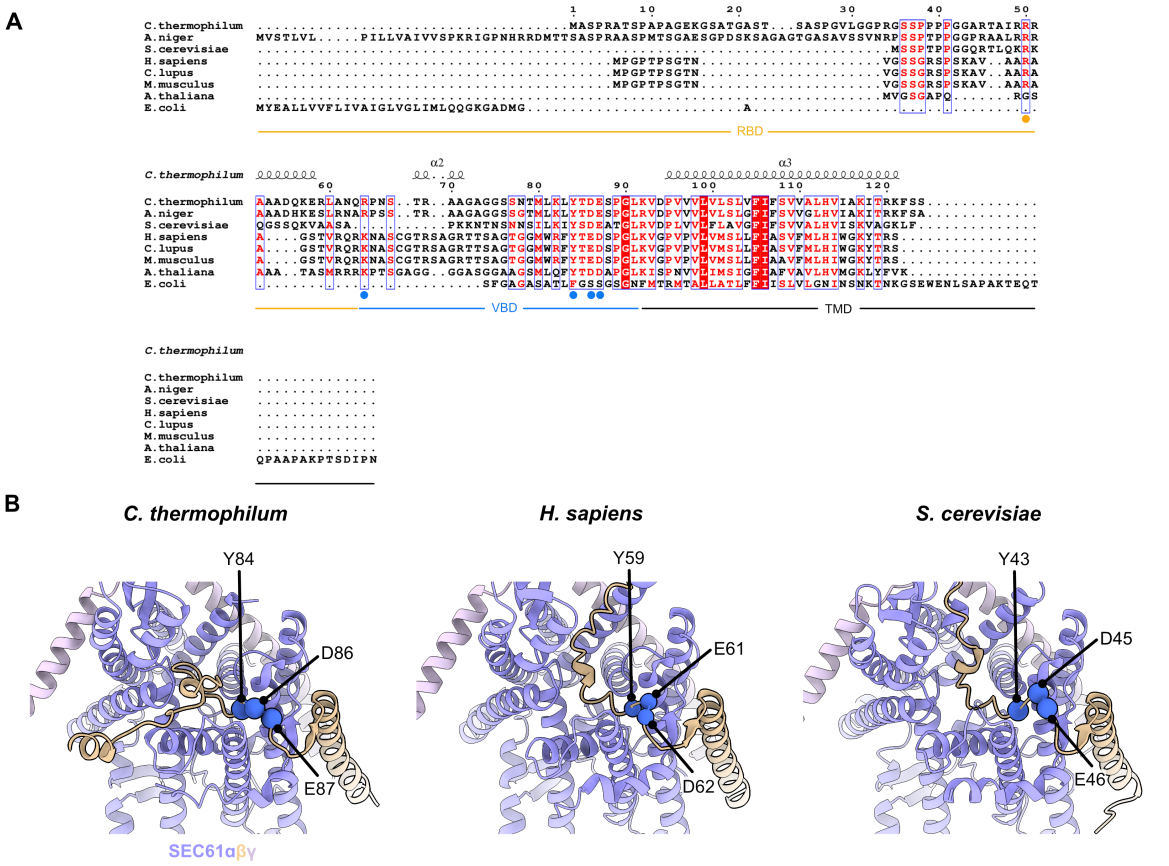


**Supplementary Fig. 14. The interaction between the SEC61β vestibule binding domain and SEC61α is conserved.**

(**A**) Sequence alignment of SEC61β generated using Clustal Omega[87] and visualised using Espript 3.0[88]. Circles below the alignment highlight conserved residues involved in interactions of the ribosome binding domain (RBD; orange) and vestibule binding domain (VBD; blue) with the ribosome and SEC61α respectively. (**B**) Side-by-side comparison of the *ct*SEC61 translocon structure and AF3[22] models for the *H. sapiens* (RMSD 1.03 Å over 367 atoms) and *S. cerevisiae* (RMSD 0.86 Å over 236 atoms) SEC61 translocon after superimposition. The analogous interaction between the SEC61β vestibule binding domain and the SEC61α cytosolic vestibule involves three of the highly conserved residues from SEC61β in **A**, shown as blue spheres.

**Supplementary Table 1. Conditions for equilibrations steps performed before production run during atomistic MD simulations.**

| | Condition | Integration time step (dt) (fs) | Simulation length (ps) | k (backbone) ($kJ\ mol^{-1}\ nm^{-2}$) | k (side chains) ($kJ\ mol^{-1}\ nm^{-2}$) | k (lipids) ($kJ\ mol^{-1}\ nm^{-2}$) | k (dihedrals) ($kJ\ mol^{-1}\ rad^{-2}$) |
|---|---|---|---|---|---|---|---|
| 1 | NVT | 1 | 125 | 4000 | 2000 | 1000 | 1000 |
| 2 | NVT | 1 | 125 | 2000 | 1000 | 400 | 400 |
| 3 | NpT | 1 | 125 | 1000 | 500 | 400 | 200 |
| 4 | NpT | 2 | 500 | 500 | 200 | 200 | 200 |
| 5 | NpT | 2 | 500 | 200 | 50 | 40 | 100 |
| 6 | NpT | 2 | 500 | 50 | 0 | 0 | 0 |
| 7 | NpT | 2 | 10,000 | 0 | 0 | 0 | 0 |

### Supplementary Videos

#### Supplementary Video 1. Structural overview of the ribosome-bound SND3 translocon.

Overview of the cryo-EM structure of the *C. thermophilum* ribosome-bound SND3 translocon. The reconstruction is colored according to the estimated local resolution values (see also **Supplementary Fig. 3**) and the model of the SND3 translocon is labeled as in **Fig. 1C**.

#### Supplementary Video 2. Lipid scrambling by SND3.

Movie of a coarse-grained (CG) molecular dynamics simulation trajectory showing the scrambling of a lipid molecule (POPC) between the membrane leaflets. SND3 and SEC61a are shown as red and blue surfaces respectively. The transparent yellow spheres represent the PO4 CG beads in the lipid molecules (POPC, POPI, and POPE) and provide reference for the lipid bilayer.
